# Human-Mediated Dispersal and Breeding Reshape Global Genomic Patterns in Black Soldier Flies

**DOI:** 10.1101/2025.08.19.670820

**Authors:** Peter Muchina, Johnson Kinyua, Fathiya Khamis, Chrysantus M. Tanga, Rawlynce Bett, Geoffrey Ssepuuya, Dorothy Nakimbugwe, Mikkel-Holger S. Sinding, Grum Gebreyesus, Goutam Sahana, Zexi Cai

**Affiliations:** Department of Biochemistry, Jomo Kenyatta University of Agriculture and Technology (JKUAT), Nairobi, Kenya; Center for Quantitative Genetics and Genomics, Aarhus University, Denmark; International Centre of Insect Physiology and Ecology (ICIPE), Nairobi, Kenya; Department of Animal Production, University of Nairobi, Kenya; Department of Food Science and Technology, Kyambogo University, Uganda; Department of Food Technology and Nutrition, Makerere University, Uganda; Department of Biology, University of Copenhagen, Denmark

**Keywords:** Black soldier fly, Whole-genome sequencing, Population genomics, Genetic diversity, Inbreeding, Founder effects, Sustainable insect farming

## Abstract

Human activities, either intentional or unintentional, have significantly influenced the global distribution and genetic composition of many species. Black soldier fly (*Hermetia illucens*; BSF) is a species that has rapidly gained commercial importance due to its bioconversion efficiency of upcycling organic waste into new products of higher value and quality through a circular economy approach. Despite its global distribution, the current Old-World demography of the wild and captive BSF populations remains poorly understood. This work combined whole-genome sequencing and population genomic analyses to determine the genetic diversity, population structure, and historical spread of global wild and captive BSF populations. Our results reveal that most global captive BSF lines were largely derived from a single primary captive lineage, likely from North America. In contrast to the genetically diverse and geographically structured wild populations, captive populations consistently exhibited reduced heterozygosity, elevated inbreeding, and extensive runs of homozygosity. These patterns reflect demographic processes such as founder effects and genetic drift, rather than intentional selection or domestication. This strongly highlights the lasting genomic impact of human-mediated dispersal and uncoordinated breeding practices. Thus, there is an urgent need for genetically informed management strategies to ensure long-term viability, adaptability, and productivity of BSF for sustainable organic waste bioconversion.

## Introduction

The black soldier fly (*Hermetia illucens*; BSF) is increasingly recognized as a promising organic waste management engineer worldwide ^1–6^. Its remarkable potential in bioconversion of organic waste into nutrient-rich larval biomass for sustainable animal feed production, frass fertilizer, and reduction of environmental pollution has accelerated their adoption into commercial production systems ^7–14^. However, despite rapid global commercialization, the genetic origins and impacts of human-mediated introduction on BSF populations remain poorly understood.

Historical records suggest that BSF, originally native to the Americas, was introduced into Africa and Europe before the widespread establishment of global captive colonies ^15^. It has been proposed that most contemporary captive populations trace their lineage back to a single or limited set of founder populations, primarily from North America ^16^. This scenario raises critical concerns about genetic bottlenecks, inbreeding, and diversity loss, which could threaten the sustainability and productivity of BSF farming practices ^17,18^.

Whole-genome sequencing (WGS) and population genomic tools provide a powerful approach to investigating genetic diversity, population structure, and evolutionary history across model and non-model organisms ^19–22^. These technologies enable high-resolution analysis of genome-wide variation, even in species with limited prior genomic resources. Although deep sequencing costs remain relatively high, cost-effective alternatives such as low-coverage WGS followed by genotype imputation allow for large-scale population analyses at reduced cost ^23–26^. These approaches are particularly valuable for emerging bio-industrial species like that of BSF, where population genetics remain largely uncharacterized and formal breeding programs are still under development.

Previous studies showed that BSF exhibits substantial genetic diversity and a structured global genetic architecture, with unique regional gene pools identified worldwide ^16,27–32^. However, comprehensive, genome-wide evidence detailing the effects of human-mediated introductions, captive breeding, and genetic exchanges between wild and captive populations remains limited ^17,18,33,34^. Moreover, limited information exists on the genetic composition of mass-reared colonies, especially in East Africa, where the BSF industry is rapidly scaling. To address this knowledge gap, we conducted WGS of 249 BSF individuals sampled from wild and captive populations in Kenya and Uganda. This was integrated with 101 globally sourced genomes from public repositories. Our goals were to: (1) characterize genome-wide patterns of genetic diversity and structure; (2) infer demographic processes shaping wild and captive lineages; and (3) assess the impact of human-mediated dispersal and breeding on global BSF population genetics.

## 2. Methods

### 2.1 Sample Collection and Sequencing

A total of 249 BSF samples (69 wild and 180 captives) were collected from different agroecological zones in Kenya and Uganda (Supplementary Data 1). Wild specimens were sampled directly from natural breeding habitats, while captive individuals were obtained from commercial and research farms, as well as smallholder producers, to capture management-driven genetic variability. DNA was extracted from all the samples using the Isolate II Genomic Extraction kit (Bioline, London, UK) following the manufacturer’s instructions. Purity and concentration were assessed using a nanodrop 2000/2000c spectrophotometer (Thermo Fischer Scientific, Wilmington, USA), while integrity was evaluated using Agarose Gel Electrophoresis (Concentration of Agarose Gel: 1%, Voltage: 100 V, Electrophoresis Time: 60 min). WGS was performed using the BGI DNBSEQ platform. The 69 wild BSF larvae were sequenced at 10× depth, while the 180 captive larvae were sequenced at 1× depth. Sequencing reads were processed using SOAPnuke ^35^ with the parameters -n 0.001 -l 10 -q 0.5 --adaMR 0.25 --polyX 50 -- minReadLen 150. The filtering procedure involved removing reads that contained 25% or more adapter sequences, permitting up to two mismatches. Reads shorter than 150 bp were discarded, while those with N content exceeding 0.1% were also removed. Additionally, sequences with homopolymer stretches (A, T, G, or C) longer than 50 bp were filtered out. Lastly, reads were excluded if at least 50% of their bases had a quality score below 10.

### 2.2 Variant Calling, Reference Panel Construction, and Imputation

An additional 101 (19 wild and 82 captive) publicly available (as of April 2024) high-coverage WGS BSF samples were downloaded from the European Nucleotide Archive (ENA) (<u>PRJNA917807</u>, <u>PRJEB58720</u>, <u>PRJEB37575</u>, <u>PRJNA196337</u>) ^27,33,36^ (Supplementary Data 2). Reads were then mapped to the BSF reference genome, <u>GCA_905115235.1</u> ^33^ using BWA-MEM v0.7.17 ^37^, and variant calling was subsequently performed using Freebayes v1.3.7 ^38^. Variant calling was performed on all samples available at the time (169 samples), excluding 180 captive samples sequenced at 1×. These samples were used as reference samples in this study. We restricted calls to biallelic sites using BCFtools v1.11 ^39^ and subsequently filtered for quality, missingness, and depth. See (Supplementary information) for additional details.

The final variant call file (VCF), filtered to exclude chromosome 7 (sex chromosome), was phased using Beagle 5.1 ^40^ with no imputation (impute=false) and tags filled using BCFtools + fill-tags ^39^. To validate the imputation accuracy of the reference panel, we randomly selected 33 high-coverage samples and downsampled them to 0.5×, 1×, and 3× coverage using SAMtools ^39^. These samples were excluded from the phased panel before imputation using QUILT v1.0.5 ^41^. Imputation accuracy was assessed by comparing imputed genotypes to the original high-coverage genotypes using the squared correlation coefficient (R²) implemented in VcfppR ^42^. Following validation, the reference panel was used to impute variants for 180 captive samples sequenced at 1× coverage. The resulting imputed VCF was merged with the reference panel using BCFtools merge. For these samples, where “true” genotypes were unavailable, the INFO score reported by QUILT v1.0.5 was used to assess the quality of imputed genotypes.

### 2.3 Population Structure Analysis

SNPs with a minor allele frequency (MAF) > 0.05 were retained using VCFtools v0.1.17 ^43^, and linkage disequilibrium (LD) pruning was performed using PLINK v1.90p PLINK v1.90p ^44^ with the parameters *--indep-pairwise 50 10 0.2* for population structure analysis. To analyze the phylogenetic relationships, the VCF file containing the population variation information was converted into a PHY file using TASSEL v5 ^45^. A maximum likelihood (ML) tree with *Ptecticus aurifer* as the outgroup was constructed by IQ-TREE v2.1.4 ^46^. The reliability of the model ML tree was estimated using the ultrafast bootstrap (UFboot) method with 1000 repeats, and the best-fit model TVM+F+I+R8 was used as the evolutionary mutation model to build the tree. The tree was visualized using Interactive Tree Of Life (iTOL) v7 ^47^. The same dataset was employed for principal component analysis (PCA) using PLINK v1.90p based on the variance-standardized relationship matrix ^44^. The first three eigenvectors were retained to create a plot in two dimensions by the R package ggplot2. We inferred the population structure by ADMIXTURE v1.3.0 ^48^, with the number of clusters (K) set from 2 to 10. The R package Pophelper v2.3.1 ^49^ was used to generate a stacked distribution bar diagram. To further explore historical splitting and admixture among populations, the same dataset with no missing values was used to build a tree using TreeMix v1.13 ^50^.

### 2.4 Genetic Diversity and Population Differentiation

According to the clustering results, genetic differentiation (F*_ST_*) and sequence divergence (D*_XY_*) were estimated using an LD-pruned VCF, with non-overlapping 50 kb windows analyzed at 50 kb intervals, each containing at least 50 genotyped sites (-w 50000 –m 50 –s 50000), implemented in popgenWindows.py (https://github.com/simonhmartin/genomics_general). Nucleotide diversity (π) was calculated using the unpruned VCF in non-overlapping 100 kb windows, with a minimum of 100 genotyped sites per window (-w 100000 –m 100 –s 100000), also using popgenWindows.py. Tajima’s *D* was computed with VCFtools using a 100 kb sliding window approach.

### 2.5 Heterozygosity

Individual-level heterozygosity was estimated in ANGSD ^51^ using individual-level site frequency spectra, measured as the proportion of heterozygous loci per sample. The SAMtools genotype likelihood model was utilized in ANGSD (-GL 1), with minimum quality filters on base quality (-minQ 20), while reducing the amount of reads with excessive mismatches (-C 50).

### 2.6 Runs of Homozygosity and Inbreeding

Runs of homozygosity (ROH) were identified using PLINK v1.90p ^44^ with the *--homozyg* function to detect ROH regions. The analysis was performed separately for each population, using only bi-allelic variants segregating within that population. A minimum ROH length threshold of 1 Mb was applied, allowing up to three heterozygous sites per window *(--homozyg-window-het 3*) and up to 20 missing genotypes per window *(--homozyg-window-missing 20*). The inbreeding coefficient ^52^ based on ROH (F_ROH_) was calculated as the total length of ROH segments (L_ROH_) divided by the total length of the autosomal genome (L).

### 2.7 Relatedness Analysis

Pairwise genetic relatedness within each population was estimated using KING v2.2.7 ^53^ with the *--related* algorithm, which calculates kinship coefficients between individuals. This analysis aimed to quantify within-population relatedness in captive and wild groups, providing context for observed patterns of inbreeding, genetic drift, and linkage disequilibrium.

### 2.8 Linkage Disequilibrium (LD) and Effective Population Size (Ne)

The linkage disequilibrium (LD) was calculated using PopLDdecay v3.43 ^54^, limiting the analysis to variants within 500 kb of each other (**-**MaxDist 500) and excluding variants with MAF below 5% (**-**MAF 0.05). The variant sets for the captive and wild (Kenya and Uganda) populations were used to estimate the effective population size (Ne) using SNeP v1.1 ^55^, with default parameters.

## Results

### 3.1 Sampling, Filtering and Imputation Accuracy

A total of 249 BSF samples were analyzed, comprising 69 wild and 180 captive individuals collected across diverse agroecological zones in Kenya and Uganda (Fig. 1B). Additionally, 101 publicly available BSF genome samples from ENA were incorporated to expand geographic representation and improve the robustness of genomic analyses (Fig. 1A). Following variant calling and quality filtering, the final dataset contained 29,894,019 biallelic SNPs, distributed across seven chromosomes (Supplementary Data 3).

**Fig 1:**
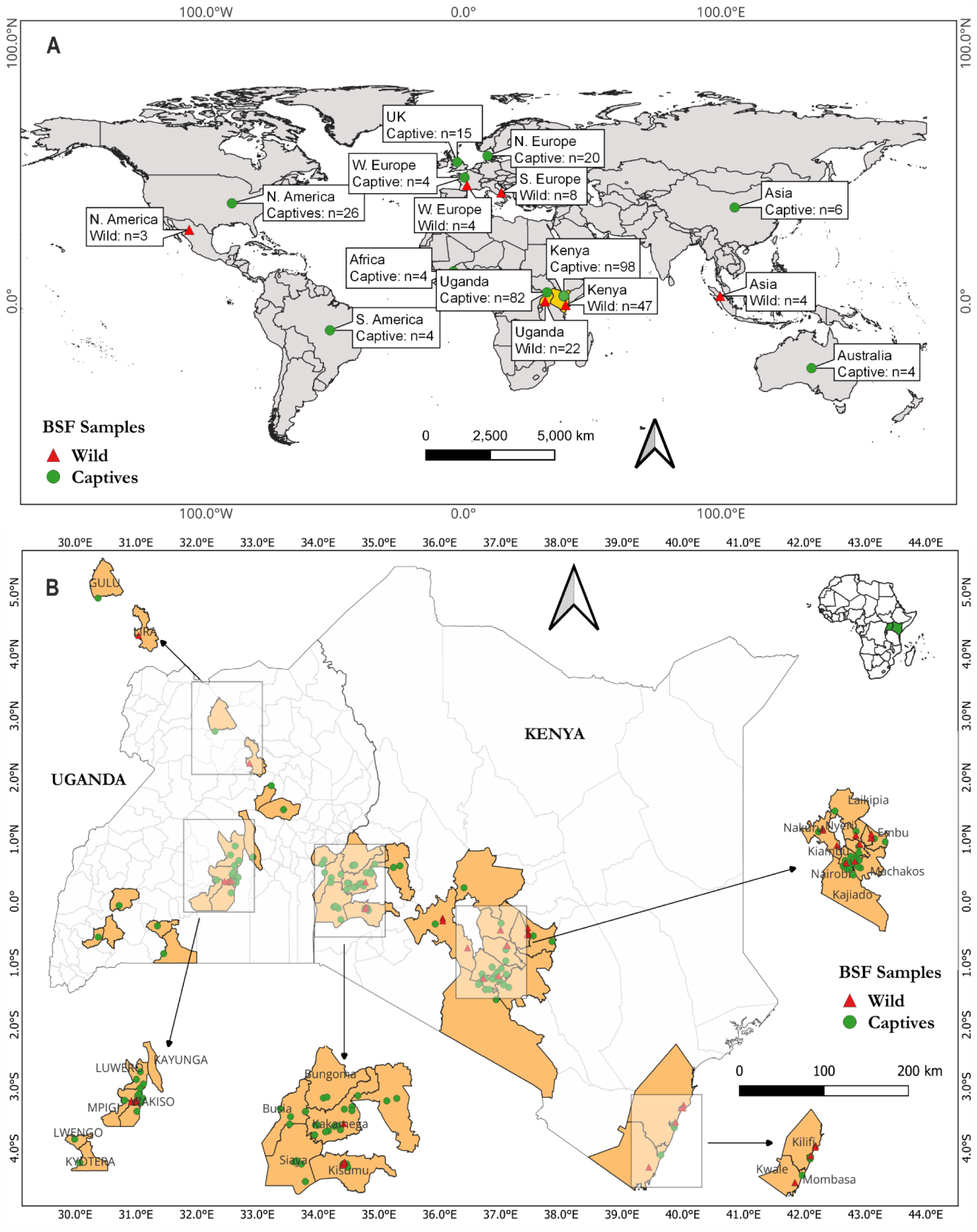
Global and Regional Distribution of Black Soldier Fly (BSF) Populations. **A,** Global distribution of wild (red triangles) and captive (green circles) BSF populations. Numbers within the boxes indicate sampled individuals for each location. **B,** Detailed sampling sites across Kenya and Uganda, showing the spatial extent of sampling efforts. Captive sites (green circles) and wild sites (red triangles) are shown.

To increase genotyping accuracy in the 1× captive samples, we applied genotype imputation using a phased reference panel constructed from high-coverage samples. To evaluate imputation accuracy, 33 high-coverage samples were downsampled to 0.5×, 1×, and 3× and imputed using the same reference panel. The R² between imputed and true genotypes improved with sequencing depth: 0.79 at 0.5×, 0.81 at 1×, and 0.84 at 3× (Supplementary Data 4, Supplementary Fig. 1). We then applied the same imputation pipeline to the full set of 180 low-coverage captive samples. INFO scores generated by QUILT v1.0.5 indicated consistently high imputation quality across these samples, with a mean INFO score of 0.92 (Supplementary Fig. 2), supporting the reliability of downstream population genomic analyses.

### 3.2 Independent Wild BSF Introductions Preceding Captive Establishment

Principal component analysis (PCA; Fig. 2A) reveals distinct clusters corresponding to wild populations from Kenya, Uganda, Europe, and Asia, each maintaining genetic identities separate from captive populations. Notably, wild populations from Kenya and Uganda cluster closely together, distinct from European and Asian wild populations. Phylogenetic reconstruction (Fig. 2B) showed well-supported and geographically distinct clades. The African population (Kenya and Uganda) forms a cohesive clade separate from European and Asian wild populations. ADMIXTURE analysis at K=4 (Fig. 2C), which had the lowest cross-validation error (Supplementary Fig. 3), showed that each wild population primarily belonged to a unique genetic cluster. Admixture levels were low across wild populations, indicating limited genetic exchange. A subset of captive samples from South America, West Africa, and Australia clustered closely with wild populations in PCA and phylogenetic analyses.

**Fig 2:**
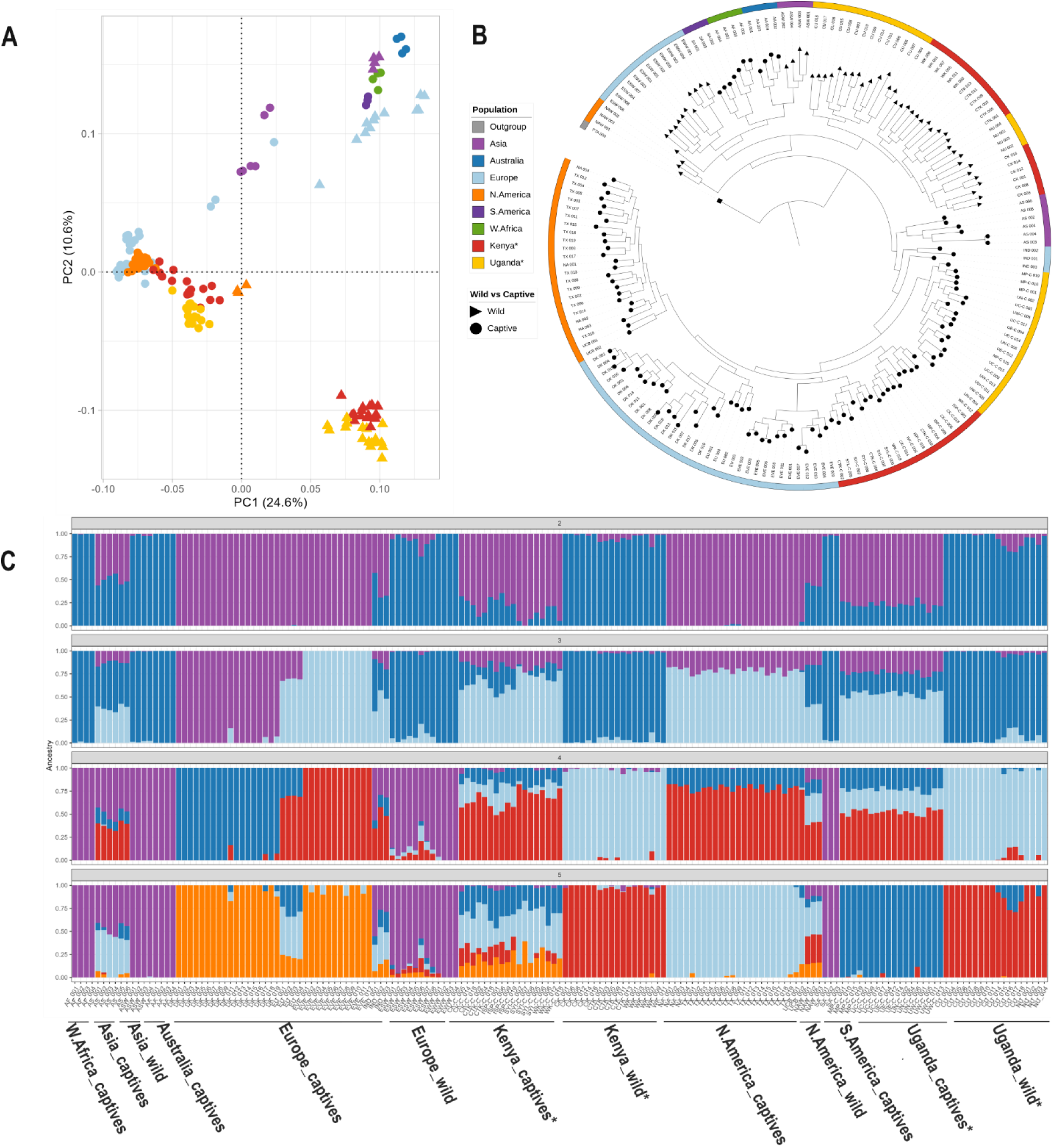
Population structure and divergence among global Black Soldier Fly Populations. **A,** PCA plot of the first two components with SNPs, with different colors representing different populations. **B,** ML tree based on SNPs with *Ptecticus aurifer* (PA) as the outgroup. **C,** Admixture based on SNPs (K=2–5). The colors in each column represent the proportion of individual genomes in each ancestral population

### 3.3 The Current Global Captive BSF Lines Originate from a Single Population

It has been hypothesized that the majority of the global BSF production lines are derived from a single line ^16^. TreeMix analysis (Fig. 3), PCA, and Phylogenetic tree (Fig. 2a and 2b), revealed a compelling signal that the majority of global BSF captive populations trace back to a single ancestral origin. Captive populations from Kenya, Uganda, North America, Europe, and Asia clustered together, diverging from a single internal node with minimal drift among them. In contrast, captive populations from South America, West Africa, and Australia fell outside this core cluster but grouped closely with the wild populations.

**Fig 3:**
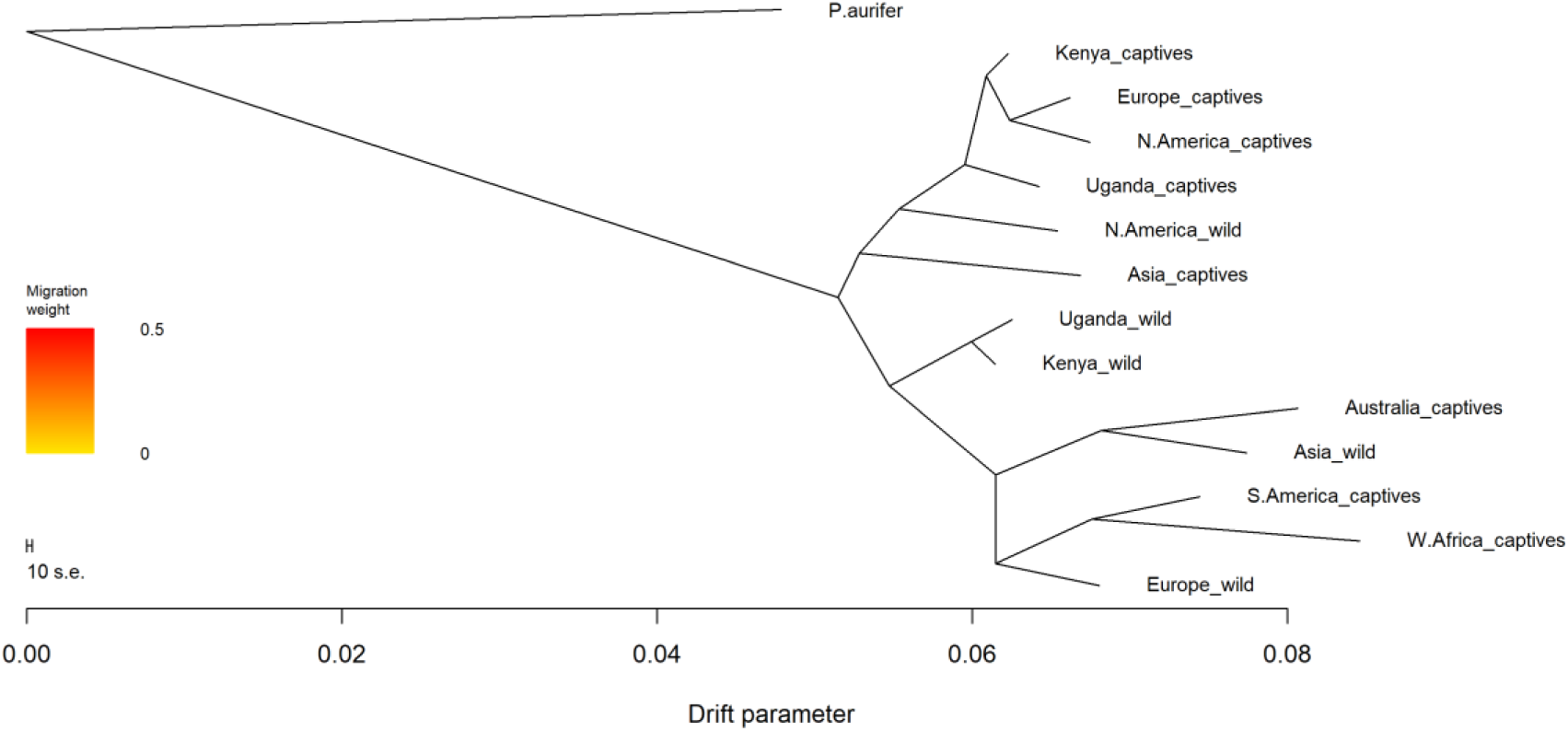
TreeMix analysis of Black Soldier Fly Populations. TreeMix reveals a shared origin for global captive populations and genetic divergence driven by drift and human-mediated dispersal. *Ptecticus aurifer* was used as the outgroup.

### 3.4 Breeding Progress is Mostly Captivity Instead of Domestication

This work assessed the genetic divergence, differentiation, and allele frequency spectra across populations using pairwise F_ST_, D_XY_, and Tajima’s *D* metrics. Pairwise F_ST_ values among many of captive populations were consistently low (e.g., < 0.10 between Kenya_captive, Europe_captive, and N. America_captive populations), indicating little genetic differentiation. Similarly, D_XY_ values among these groups were also low, suggesting limited nucleotide divergence and supporting a model of recent shared ancestry with minimal evolutionary distance. In contrast, comparisons between wild and captive populations showed slightly elevated F_ST_ values, yet these remained modest (e.g., ∼0.05–0.10), far below what would be expected if strong, long-term selection had shaped captive genomes. Tajima’s *D* values (Fig. 4B) were generally positive for most captive populations, including those from Asia, Europe, and North America. In contrast, wild populations had values near zero or slightly negative.

**Fig 1:**
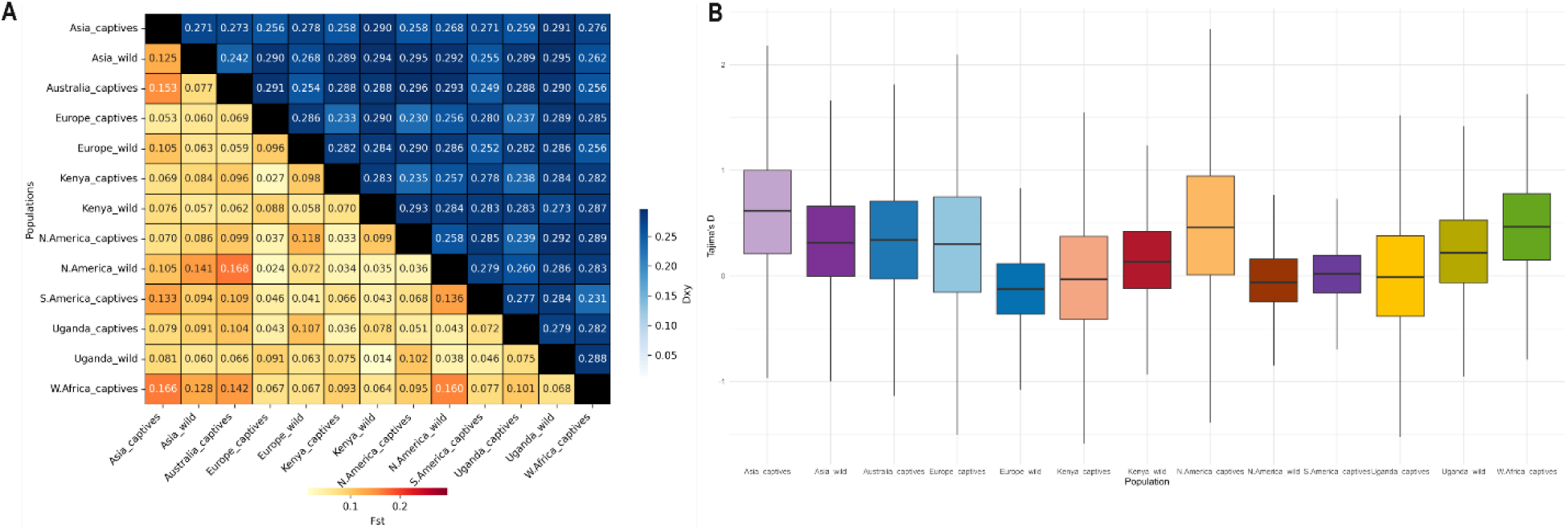
Genetic differentiation and neutrality test across Black Soldier Fly Populations. **A,** Genetic differentiation as described by F_ST_ (yellow-red) and D_XY_ (blue) for black soldier fly populations, rounded to three significant figures. **B,** Tajima’s *D* across different black soldier fly populations.

### 3.5 The Main Force Shaping the Current Captive Lines

Linkage disequilibrium decay curves (Fig. 5A) clearly distinguish between captive and wild populations. Wild populations from Kenya, Uganda, and Europe exhibit rapid LD decay, consistent with large effective population sizes and frequent recombination. In contrast, most captive populations maintain elevated LD across long genomic distances. Notably, recently established captive lines from South America, West Africa, and Australia also displayed elevated LD.

**Fig. 5:**
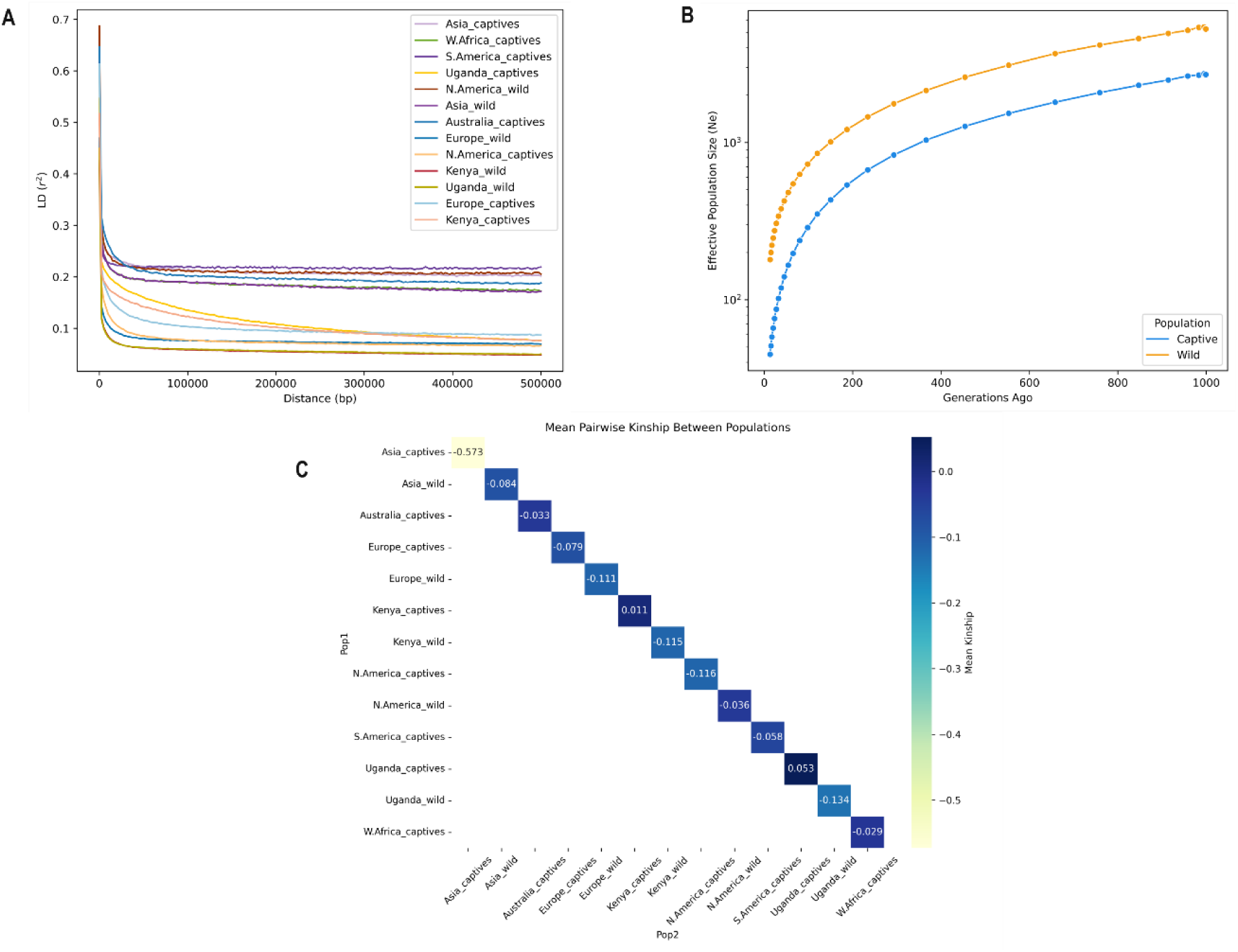
Genomic signatures of demographic change across black soldier fly populations. **A,** Linkage disequilibrium (LD) decay across genomic distance, contrasting wild and captive populations. **B,** Historical effective population size (Ne) estimates derived from genome-wide SNP data. **C,** Pairwise genetic relatedness within populations, highlighting variation in relatedness across captive and wild groups.

Estimates of historical effective population size (Ne) (Fig. 5B) show a marked reduction in captive populations relative to their wild counterparts. Kinship coefficients (Fig. 5C) revealed elevated values within populations in some captive groups, particularly in Kenya and Uganda, consistent with reduced diversity and close relatedness. In contrast, wild populations display uniformly negative kinship values. Notably, the Asia_captive population displayed the exceptionally negative mean kinship (–0.573) relative to all other populations and also exhibited the highest within-population mean kinship (0.2455) (Fig. 5C; Supplementary Data 5). Network visualization of individual kinship relationships (Supplementary Fig. 5) illustrates dense clustering among captive colonies primarily from Kenya, Uganda, and Europe. These patterns distinguish captive populations from their wild counterparts and indicate that common demographic signals, such as elevated linkage disequilibrium, reduced effective population sizes, and increased relatedness, consistently characterize the current global captive lines.

### 3.6 Severe Inbreeding in Captive BSF Populations

The genetic consequences of captive BSF breeding were examined through heterozygosity, runs of homozygosity (ROH), and inbreeding coefficients (F_ROH_) across populations. Compared to their wild counterparts, the observed heterozygosity (Fig. 6A) was significantly lower in most captive populations, especially from Asia, Europe, Kenya, and Uganda. Total ROH length (Fig. 6B) was substantially higher in captive populations, particularly from Asia and Europe. The elevated ROH in the Asia BSF population coincides with highly negative kinship values, while the Europe captive population showed tightly clustered individual relationships (Supplementary Fig. 5). The inbreeding coefficient F_ROH_ (Fig. 6C) further quantified these patterns, and the Asia_captives exhibited the highest F_ROH_, followed by Europe_captives and Kenya_captives. In contrast, wild populations showed near-zero ROH and F_ROH_ values.

**Fig 6:**
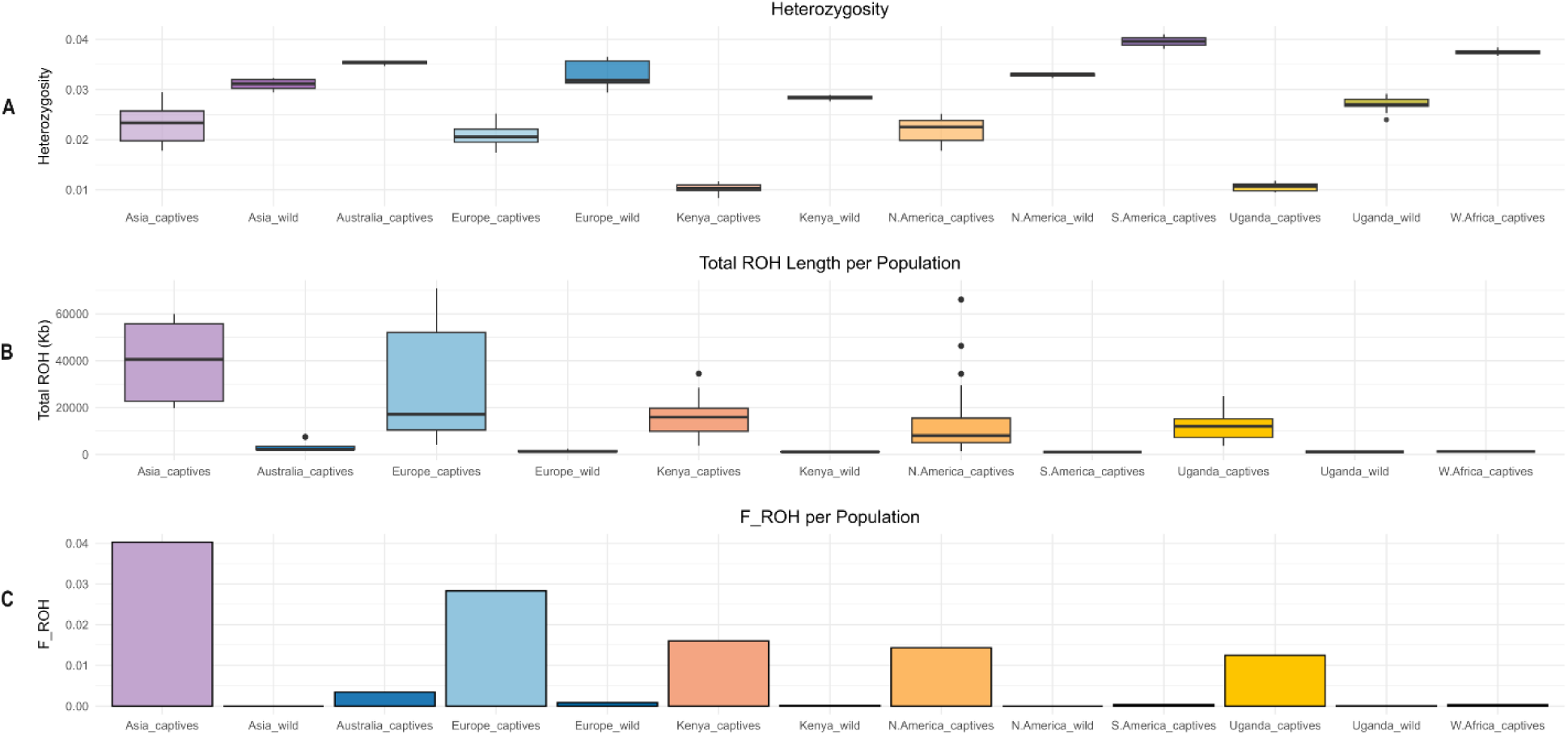
Genetic diversity and inbreeding patterns across BSF populations. **A**, Genome-wide heterozygosity. **B**, Total length of runs of homozygosity (ROH). **C**, Inbreeding coefficient (F_ROH_) estimated from ROH. Captive populations show variable reductions in diversity and elevated inbreeding compared to wild populations.

### 3.7 Summary of Demographic Forces Across Populations

We integrated results from TreeMix phylogenetic inference, genome-wide heterozygosity, ROH, F_ROH_, LD decay, Ne, and pairwise relatedness to summarize the demographic forces shaping each BSF population. Core captive populations, particularly in Asia and Europe, exhibited extensive ROH segments, high F_ROH_, slow LD decay, and elevated relatedness consistent with strong genetic drift and inbreeding. In contrast, wild populations from Kenya, Uganda, Europe, and Asia showed high heterozygosity, short ROH, low F_ROH_, and rapid LD decay, indicative of large, outbred populations with limited drift. Captive populations in South America, West Africa, and Australia maintained high diversity, with LD patterns resembling wild lineages, suggesting recent establishment from wild sources. The integrated synthesis map (Fig. 7B) illustrates these regional differences and highlights the contrasting demographic histories that underline global BSF genomic structure.

**Fig 7:**
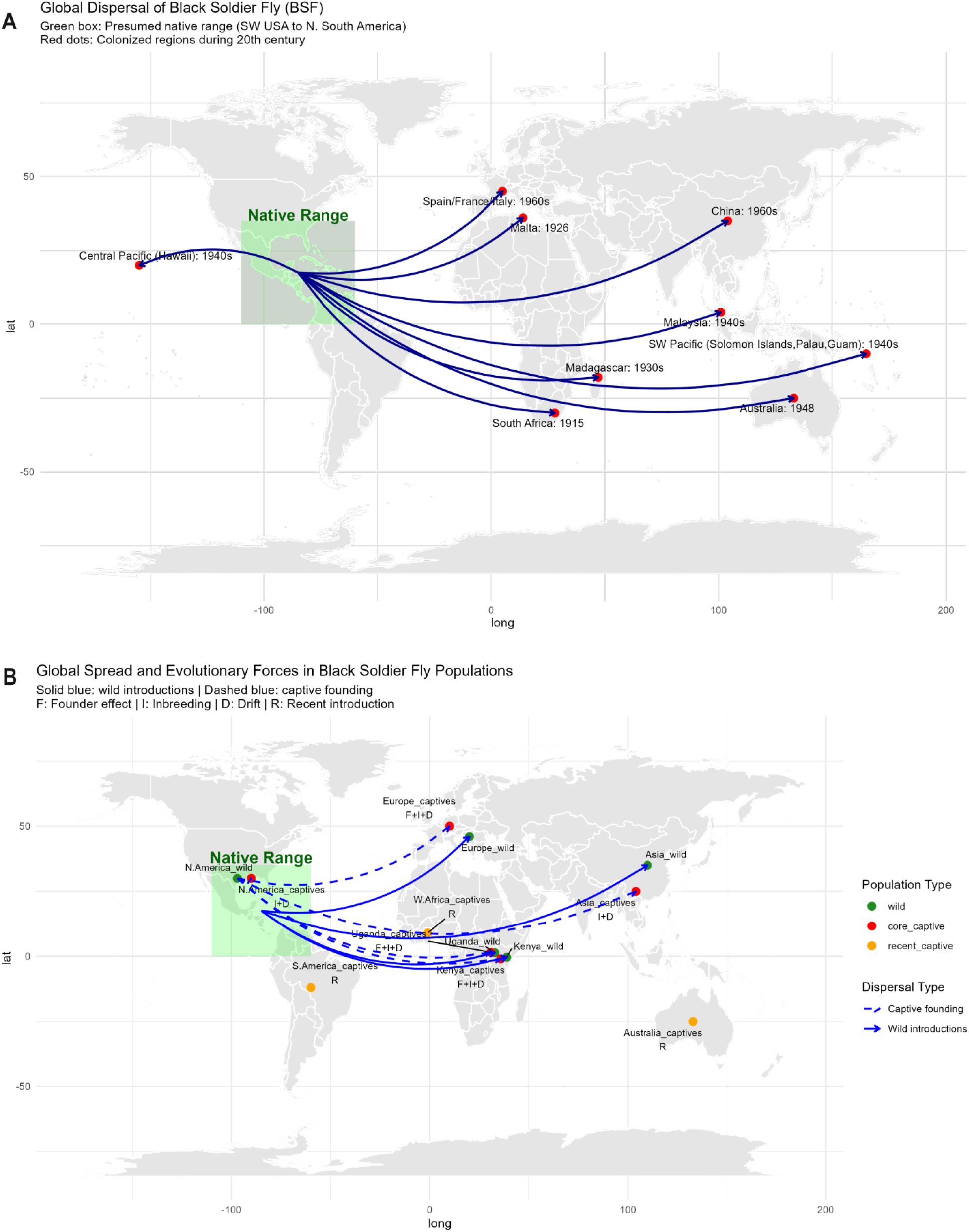
Global dispersal and demographic history of the black soldier fly (*Hermetia illucens*). **A**, Historical dispersal inferred from records of global spread during the 20th century. The shaded green box denotes the presumed native range (southwestern USA to northern South America). Arrows indicate introductions to other continents; red points mark documented introduction sites with approximate dates. **B**, Genomic synthesis of dispersal and demographic forces. Solid blue arrows represent introductions from the native range to wild populations; dashed blue arrows indicate founding of captive lines from North American wild flies. Circle colors denote population categories (green = wild, red = core captive, orange = recent captive). Population labels include dominant evolutionary forces inferred from genomic data: F = founder effects, I = inbreeding, D = genetic drift.

## Discussion

This study leveraged WGS to investigate the global population structure of BSF, focusing on wild and captive populations. Our finding reveals that most captive lines across continents trace their ancestry to a single population, while wild populations remain genetically distinct and geographically structured. These findings underscore the dominant role of human-mediated dispersal and drift in shaping captive BSF genomes and highlight the importance of preserving wild diversity to support sustainable breeding and long-term productivity.

### A Global Industry Rooted in a Single Ancestral Line

Genomic analyses provide strong evidence that the majority of captive populations across Africa, Europe, Asia, and the Americas trace their ancestry to a single domesticated lineage ^16^ (Fig. 3). Phylogenetic reconstruction, PCA, ADMIXTURE, and TreeMix analyses consistently revealed tight clustering of captive populations with low levels of drift. This pattern supports a model of centralized domestication, likely originating in North America, a recognized hub for early BSF research and commercial breeding ^16^. This centralized origin is further supported by the remarkably low pairwise F_ST_ and D_XY_ values observed among global captive populations, indicating recent shared ancestry and minimal evolutionary divergence ^56,57^. Such genetic uniformity is consistent with the rapid, human-mediated dissemination of a founding stock to meet growing industrial demand for BSF in organic waste conversion and alternative protein production ^12,58^.

However, several exceptions to this pattern were observed. Captive populations from Australia, South America, and West Africa clustered closely with their respective wild populations, a pattern also supported by their placement in TreeMix topologies. These findings, in line with previous reports ^16^ suggest that these lines were established from local wild-caught individuals, either to preserve adaptive traits or due to recent, unstructured domestication efforts. Their genetic similarity to wild counterparts reflects limited time in captivity and the absence of sustained selection or inbreeding ^18^.

### A Two-Phase Expansion Model: Natural Introduction Followed by Human-Mediated Spread

The global distribution of BSF reflects a two-step process involving early introductions through global trade, followed by more recent, human-mediated dispersal of domesticated lines. Although BSF is a cosmopolitan species in all tropical and selected subtropical and temperate regions, it is native to the Americas, ranging from Northern South America to the Southwestern United States ^15,16,28,29^. The presence of BSF in the late 19^th^ and early 20^th^ centuries has been recorded in the Americas ^15^. All the populations in Africa, Europe, and Asia are considered introduced ^28^. Historical records indicate a stepwise global expansion: first to the Afrotropics, where BSF was recorded in South Africa by 1915 and in Madagascar during the 1930s; followed by the Palearctic (Malta in 1926; Spain, France, and Italy by the 1960s), and Southeast Asia in the 1940s ^15^. In the Australian region, BSF was recorded in 1948 ^28^. In the Pacific, the species was widespread by the 1940s, coinciding with World War II shipping routes ^15,28^. These patterns, with early records from the islands or coastal hubs, strongly support worldwide trade as a major route for its spread (Fig. 7A).

Our genomic analyses align with this dispersal history. Phylogenetic and PCA analyses show that wild BSF populations in Africa, Europe, and Asia are genetically distinct from each other and from the captive lines, indicating multiple independent introduction events into these regions. Kenyan and Ugandan wild populations cluster closely together, suggesting a shared introduction event distinct from European and Asian lineages. This is consistent with early transcontinental trade and transport networks, particularly from the Americas, where BSF was first described and extensively studied ^28^. In contrast, TreeMix and ADMIXTURE results support the hypothesis that most global captive BSF populations descend from a single ancestral lineage, likely originating in North America ^16^. This supports the idea of a single domestication event followed by rapid, human-mediated dissemination through commercial rearing and colony exchange among farms and research centers. The combined evidence suggests a two-phase expansion model: (1) an accidental or intentional introduction of BSF to Africa, Europe, and Asia via global trade, followed by (2) a more recent, centralized dissemination of captive BSF lines, shaped by industrial rearing and human-assisted spread.

### Inbreeding and Genetic Drift Drive Divergence in Captivity

While “domestication” often implies intentional selection for desirable traits, it more broadly refers to a coevolutionary process in which one species manages the survival and reproduction of another to obtain resources or services, leading to long-term mutual adaptation ^59^. However, our results suggest that genetic divergence in most captive BSF populations is not primarily driven by such targeted selection. Instead, it reflects demographic processes, particularly founder effects, genetic drift, and inbreeding, stemming from unstructured colony establishment ^17,34,60^. Captive populations showed elevated linkage disequilibrium (LD), reduced effective population sizes (Ne), and consistently positive Tajima’s *D* values. These patterns are classic signatures of bottlenecks and reduced recombination in closed or recently established colonies ^17^. Recently established captive lines from South America, West Africa, and Australia displayed elevated LD, likely reflecting their limited number of generations in captivity and correspondingly few recombination events since establishment, leading to the retention of long-range LD inherited from their founders ^61^.

Further evidence of demographic-driven divergence comes from extensive runs of homozygosity (ROH), low heterozygosity, and elevated inbreeding coefficients (F_ROH_), particularly those from captive Asian and European populations. These features collectively indicate severe inbreeding due to small founding populations, restricted gene flow, and repeated use of related individuals, consistent with unstructured breeding and colony recycling ^62–65^. Kinship analyses support this view, as most captive populations exhibited higher within-group relatedness (Supplementary Data 5). The Asia_captive population displayed the highest within-population mean kinship (0.2455), reflecting a high degree of shared ancestry among individuals, consistent with sustained inbreeding and founder effects. Simultaneously, it showed a strongly negative mean pairwise kinship (–0.573) relative to all other populations (Fig. 5C), indicating substantial genome-wide divergence from the global reference panel. While negative kinship values can arise from statistical centering ^66,67^, such extreme values in a closed, bottlenecked colony often indicate prolonged demographic isolation and lack of outcrossing ^62,68^. This dual signature, high internal relatedness, and deep external divergence underscore the cumulative impact of drift, restricted gene flow, and long-term captive propagation in shaping the genetic profile of this population.

In contrast, wild populations retained high genetic diversity, low LD, near-zero F_ROH_, and large Ne estimates, indicative of panmixia, large breeding pools, and relatively stable demographic histories ^18,69,70^. Estimates of nucleotide diversity (π) were also markedly higher in wild populations, underscoring their role as critical reservoirs of genomic variation (Supplementary Fig. 4). These contrasting patterns between wild and captive populations highlight the urgent need for genetic monitoring in BSF breeding systems. Without intervention, demographic processes, rather than adaptive evolution, will continue to dominate, threatening long-term colony viability and productivity.

### Implications for Domestication and Breeding Resilience

Our findings emphasize that breeding progress in BSF has occurred under captivity, but not under structured domestication. The absence of strong divergence signals between captive populations and their wild ancestors, along with evidence of drift-driven divergence, suggests that artificial selection has not yet played a major role. This pattern is typical of early-stage insect domestication, where colonies are frequently established through opportunistic collection from wild founders without structured selection ^17,34,71,72^. Such practice introduces a population bottleneck that rarely captures the full spectrum of genetic variation and reduces effective population size from the outset. As reproductive cycles become closed, ongoing genetic drift and unintentional inbreeding further erode diversity ^17,18,73,74^. Simultaneously, natural selection is relaxed, while adaptation to artificial environments may trigger selective sweeps, compounding genetic loss ^75^. Captive BSF colonies may thus be caught in a cycle of genetic erosion, with phenotypic unpredictability, including changes in development, behavior, and robustness, that threaten colony viability and productivity ^18,72,76,77^.

The genetic homogenization observed across global industrial BSF colonies may limit their capacity to adapt to selection pressures, environmental changes, or disease outbreaks ^17,78–82^. While uniformity supports production consistency in the short term, it can hinder long-term breeding success, a pattern also observed in other r-selected insects and aquaculture species ^83–86^. These concerns are particularly relevant for regions such as East Africa, which has become a global focal point for BSF production ^87–89^. Kenya and Uganda have approved all relevant edible insect species for food and feed, in contrast to restrictive European policies ^89,90^. Moreover, over 75% of surveyed farmers and feed millers in the region have expressed readiness to adopt insect-based technologies ^91^. This combination of policy support and grassroots enthusiasm highlights East Africa’s strategic importance as an industrial hub and a reservoir of valuable genetic diversity for sustainable breeding. Our genomic analyses offer critical insights into the development of well-structured BSF breeding programs. In Kenya and Uganda, we identified significant variation in genetic diversity, inbreeding levels, and kinship among wild and captive populations. This information can directly inform the selection of optimal breeding lines or individuals. For instance, individuals from captive populations with elevated F_ROH_ and high within-colony relatedness may be poor candidates for further propagation due to the risk of inbreeding depression. In contrast, genetically diverse individuals from wild populations or recently established colonies with high effective population sizes (Ne) and low kinship values offer promising sources for expanding or restoring breeding pools.

By integrating these genomic indicators with phenotypic and production data, BSF farmers and researchers, especially in East Africa, can make data-driven decisions when selecting pedigrees, optimizing colony health, productivity, and long-term resilience. Importantly, these findings underscore the value of incorporating local wild diversity into breeding pipelines, particularly in regions like Kenya and Uganda, which serve as biodiversity hotspots and industrial production hubs for BSF.

### Future Directions

While our study includes globally representative BSF populations, certain regions, particularly in South America, remain underrepresented, possibly masking additional domestication events. The lack of complete historical records further complicates aligning genetic divergence with introduction timelines. Although our genome-wide data suggest a single major domestication origin, unsampled or under-sampled populations may reveal alternative domestication histories or novel genetic lineages.

Given the central role of BSF in sustainable protein production and organic waste management, preserving its genetic integrity is critical. Future efforts should prioritize broader geographic sampling, especially in regions with historical BSF presence or recent commercialization. Additionally, the development of high-quality genomic references and integrating functional genomics to identify traits of agronomic importance, such as growth rate, waste conversion efficiency, stress tolerance, and reproductive performance. To support breeding resilience and long-term viability, the implementation of structured, genomically informed breeding programs is urgently needed. Routine genetic monitoring to track diversity, relatedness, and inbreeding should become standard practice across BSF production systems. Such efforts will be crucial for preventing inbreeding depression, enhancing adaptability, and sustaining BSF’s role as a key bio-industrial species.

### Conclusion

This study provides compelling genomic evidence demonstrating that human-mediated activities, including the uncoordinated establishment of breeding programs, founder effects, and limited monitoring of inbreeding, have substantially influenced the global genetic structure and diversity of BSF populations. While our results support a single major origin for most captive lines, the genetic divergence across regions reflects demographic processes rather than targeted selection. This challenges conventional assumptions of domestication and underscores the importance of maintaining genetic diversity in breeding pipelines. Importantly, the wild and recently established populations retain high genetic variation and low levels of inbreeding, making them invaluable resources for sustaining future breeding efforts. Integrating genomic data into colony management can facilitate the identification of optimal pedigrees, enable ongoing monitoring of genetic health, and help prevent inbreeding depression, ultimately promoting more resilient and productive BSF farming systems. Our findings offer a global framework for managing genetic resources in this growing insect bioconversion industry and emphasize the critical need for regionally informed, genomically guided breeding programs.

## Data Availability

All data used in this study are described in the main text and Supplementary Data. Sample locations and sources are detailed in Supplementary Data 1. Raw sequencing data generated in this study and associated metadata are available from the NCBI Sequence Read Archive under BioProject accession PRJNA1256126. Publicly available whole-genome data incorporated in the analysis can be accessed under BioProject accessions <u>PRJNA917807</u>, <u>PRJEB58720</u>, <u>PRJEB37575</u>, and <u>PRJNA196337</u>. The *Hermetia illucens* reference genome assembly <u>iHerIll2.2.curated.20191125</u> is available for download from NCBI.

## Code availability

Details of all software and version numbers used in the analyses are available in the Methods section. The code used for analyses in this study is publicly available on GitHub: https://github.com/PMuchina/BSF_genomics_pipeline.

## Acknowledgements

We thank Levi Ombura and Emmanuel Kiprono from the Arthropod Pathology Unit (APU) lab, ICIPE, for their invaluable assistance with DNA extractions. We also thank Kentosse Gutu, Eric Rachami, and Mark Ojungu for supporting sample collection across Kenya and Uganda.

## Author contributions

PM: sampling, methodology, analysis, writing – original draft; JK: sampling, methodology, supervision – review & editing; FK: sampling, methodology, supervision – review & editing; CMB: sampling, methodology, supervision – review & editing; RB: sampling – review & editing; GSP: sampling – review & editing; DN: sampling – review & editing; MSS: analysis, writing – review & editing; GG: sampling, methodology, supervision – review & editing; GS: methodology, analysis, writing – review & editing; ZC: methodology, analysis, writing – review & editing. All authors proofread and approved the final version of the manuscript.

## Ethics statement

This study was reviewed and approved by the Jomo Kenyatta University of Agriculture and Technology (JKUAT) Institutional Scientific and Ethics Review Committee (ISEREC) (JKU/ISERC/02316/1107) and the National Council for Science Technology and Innovation in Kenya (NACOSTI/P/25/415352).

