## Supplementary information for "Human-Mediated Dispersal and Breeding Reshape Global Genomic Patterns in Black Soldier Flies"

*[1]Department of Biochemistry, Jomo Kenyatta University of Agriculture and Technology (JKUAT), PO Box 62000-00200, Nairobi, Kenya; [2]Center for Quantitative Genetics and Genomics, Aarhus University, C.F Møllers Alle 3, Denmark; [3]International Center of Insect Physiology and Ecology (ICIPE), PO Box 30772-00100, Nairobi, Kenya; [4]Department of Animal Production, College of Agriculture and Veterinary Sciences, University of Nairobi, P.O. BOX 29053-00625, Nairobi, Kenya; [5]Department of Food Technology and Nutrition, School of Food Technology, Nutrition & Bioengineering, Makerere University, P. O. Box 7062, Kampala, Uganda; [6]Department of Food Science and Technology, Kyambogo University, P.O. Box 1 Kyambogo, Kampala, Uganda; [7]Department of Biology, University of Copenhagen, Ole Maaløes Vej 5, 2200 Copenhagen N, Denmark*

### Supplementary Information

### VCF Filtering Steps

To ensure high-quality variant calls and improve the efficiency of downstream analyses, initial filtering was performed on the raw FreeBayes-called VCF file using three key parameters: a missingness threshold of >50%, a minor allele count (MAC) of at least 3, and a minimum site quality score (Q) of 30. These filters helped remove a substantial number of low-quality or potentially spurious variants, retaining approximately half of the original variant set and significantly improving the performance of subsequent steps. Individuals with high levels of missing data (>50%), including two samples and one outgroup, were excluded from the dataset. Block substitutions were decomposed, and duplicate variants were removed using the vt toolkit. Only biallelic SNPs were retained, as these are commonly required by most imputation tools. Additional filtering was applied based on mean depth (DP) thresholds, keeping only variants with a mean DP between 5 and 25. A stringent overall missingness threshold of 5% was enforced, followed by a population-specific missingness filter to account for variation across multiple populations. Allelic balance was assessed, and variants with extreme imbalance (outside 0.25–0.75 or below 0.01) were filtered out. To remove non-informative or monomorphic sites, a minor allele frequency (MAF) threshold of 0.001 was applied. A formatting correction was performed to address inconsistencies in allele representation. Finally, variant annotations were completed using bcftools +fill-tags to populate missing fields such as MAF, enabling further exploratory analyses. The resulting VCF file was then used for phasing and downstream analyses.

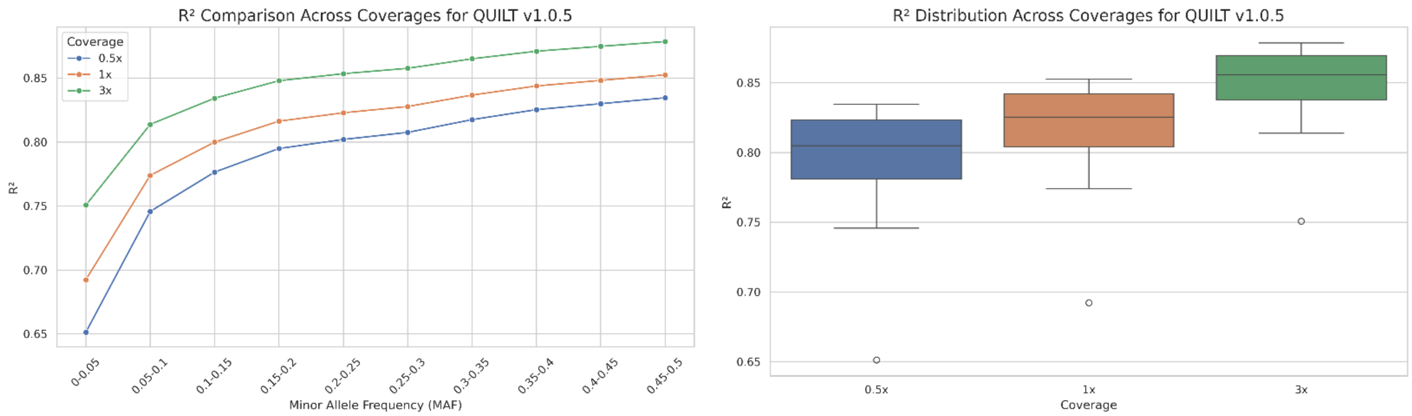

**Supplementary Figure 1: Squared correlation coefficient ($R^2$) for QUILT v1.0.5 across 0.5×, 1×, and 3× coverages.**

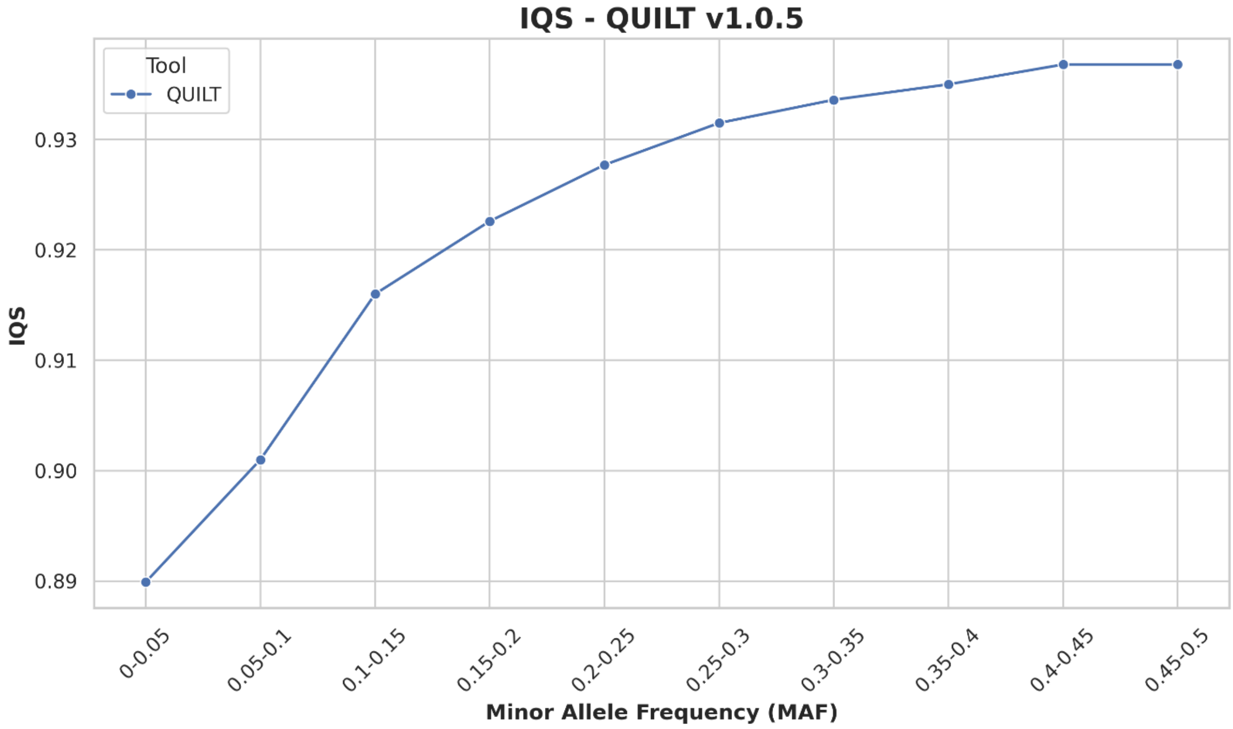

**Supplementary Figure 2: INFO scores generated by QUILT v1.0.5 for the 180 low-coverage samples (1×).**

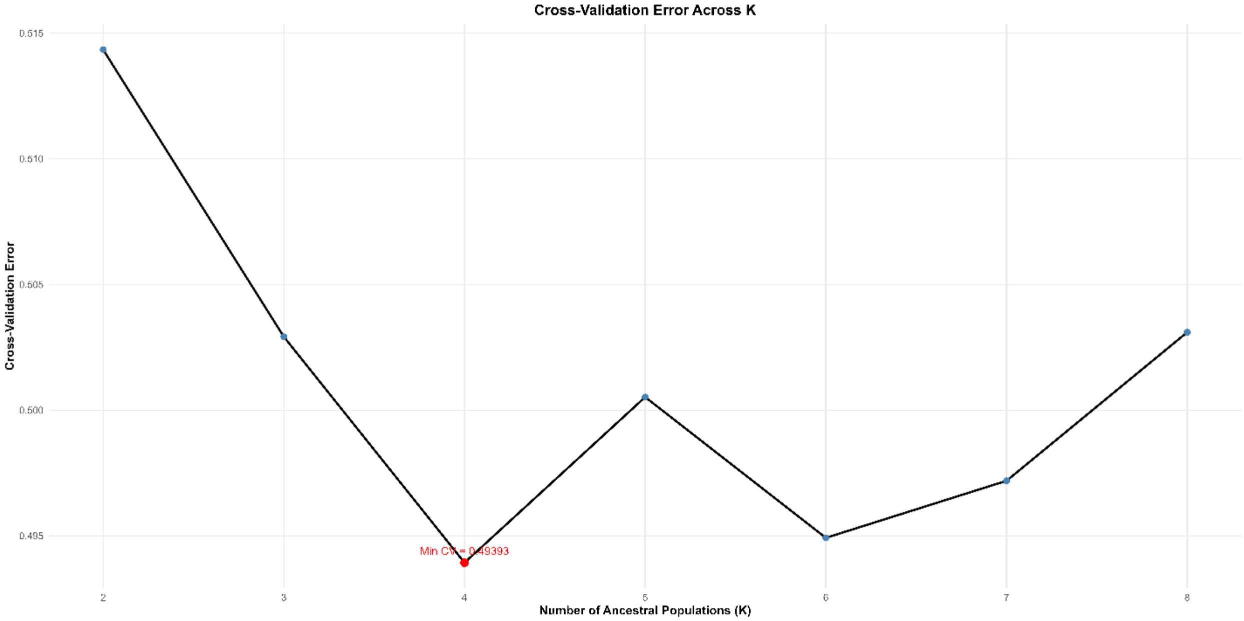

**Supplementary Figure 3: Admixture cross-validation error across K2 to K8.**

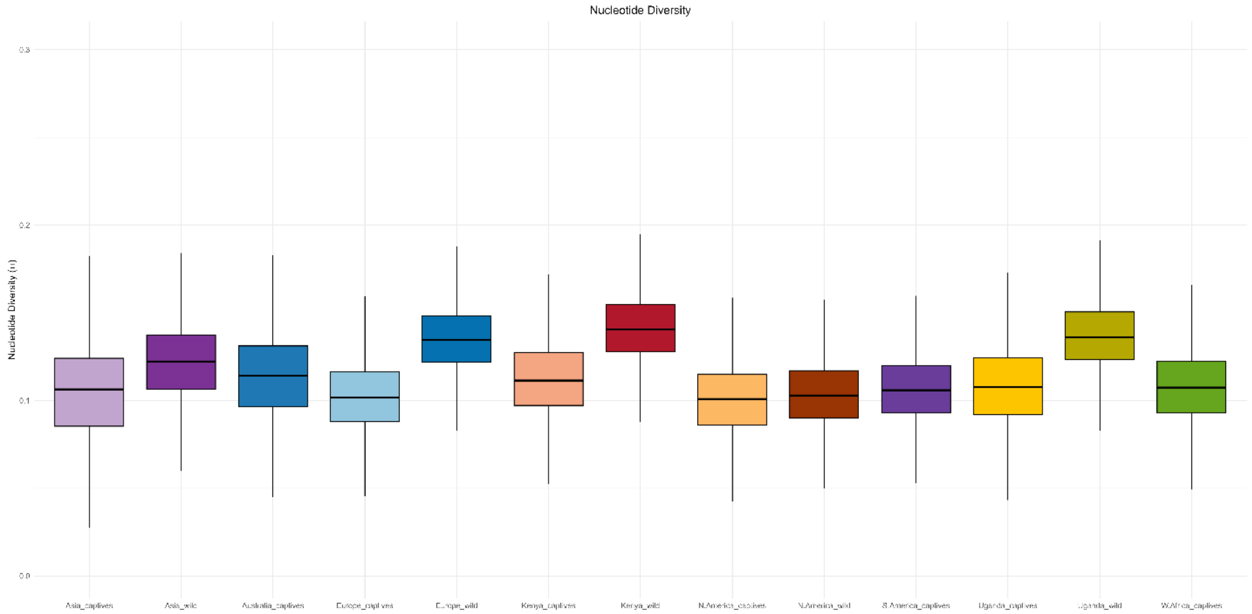

**Supplementary Figure 4: Nucleotide diversity across different black soldier fly populations.**

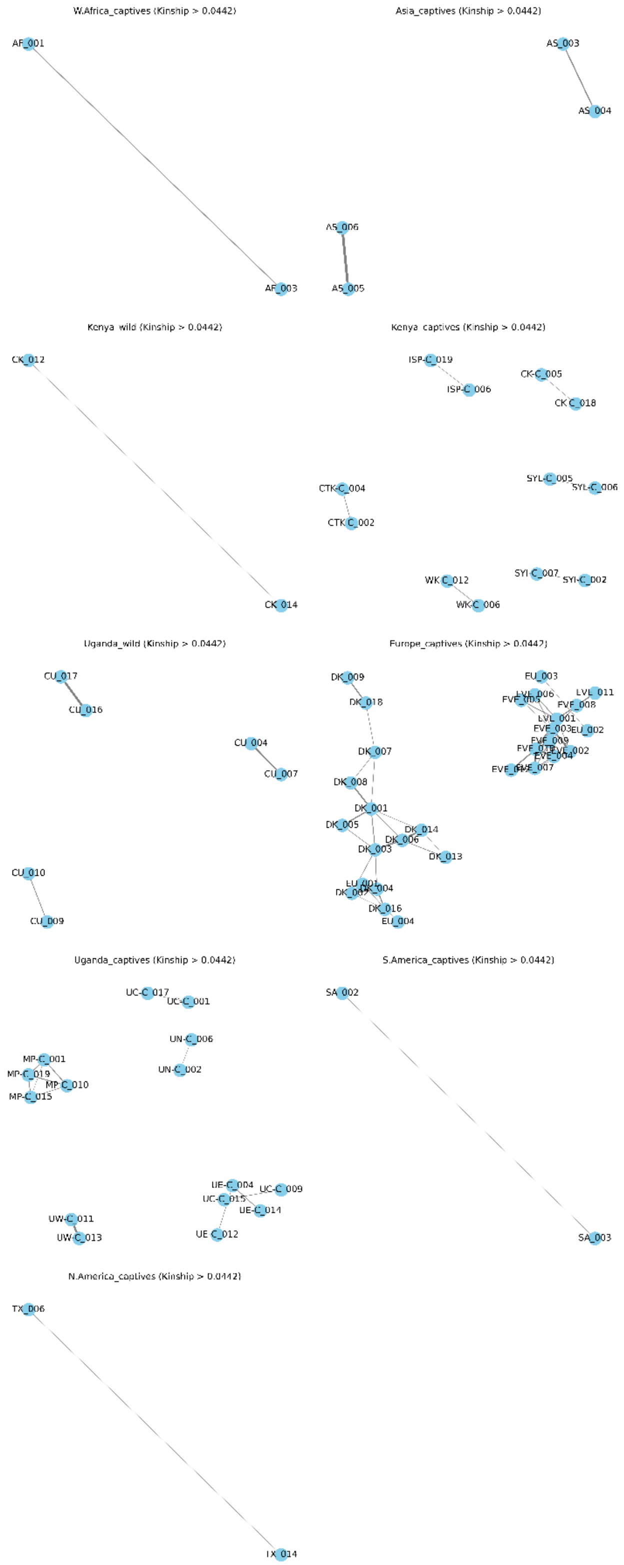

**Supplementary Figure 5: Relatedness network within global black soldier fly populations.**
